# Electroencephalographic evidence for the involvement of mirror neuron and error monitoring related processes in virtual body ownership

**DOI:** 10.1101/795773

**Authors:** Gal Raz, Guy Gurevitch, Tom Vaknin, Araz Aazamy, Iddo Gefen, Stanislaw Grunstein, Gal Azouri, Noam Goldway

## Abstract

The illusion that an artificial or virtual object becomes part of one’s body has been demonstrated and productively investigated in the past two decades. Empirical and theoretical accounts of this phenomenon suggest that the body ownership illusion relies not on a single process, but rather on the alignment of the biological and the alternative bodies across multiple aspects. However, the portrayal of these aspects and the demarcation of their neurophysiological correlates has yet to be established.

Our study examines electroencephalographic (EEG) markers of two extensively studied systems in the context of virtual body ownership illusion: the mirror neuron system (MNS) and the error monitoring system (EMS). We designed an experimental manipulation of brief involuntary virtual hand bounces, which triggers both systems, and examined how the response of EEG markers of these systems to this manipulation is modulated by three aspects of body ownership: agency, visuotactile synchronicity, and semantic congruence between the participant’s hands and its virtual representation.

We found evidence for enhanced MNS-related suppression of power at the Mu band in the synchronous and semantic congruence conditions. On the other hand, the EMS-related Pe/P300 wave was reduced by semantic congruence. This Pe/P300 effect was stronger among participants who exhibited higher acceptance of the spatial illusion and increased tendency for affective empathy. Mu power and Pe/P300 were not correlated, suggesting a dissociation between the distinct aspects of body ownership they probe. The findings suggest that synchronicity and semantic congruence induce sensorimotor sensitivity to the alternative body, whereas the latter parameter also buffers minor erroneous virtual motions. These neurophysiological markers may be added to the arsenal of body ownership probes, and integrated in VR rehabilitation protocols.

## Introduction

One of the most fundamental aspects of human consciousness is the fact that it resides in a unique biological body. An individual commonly develops a rock-solid one-to-one identification with a single body over which she feels total ownership. Intriguingly, and somewhat counter- intuitively, current research has suggested that this experience of a coherent “embodied self” actually hinges on an interplay between different processes rather than on a single mechanism (Metzinger, 2004). Among the more influential theoretical dissections of self-consciousness are Antonio Damasio’s distinction between the proto-self, core self, and autobiographical self along an axis moving from visceral and autonomic to cognitive processing (Damasio, 2012 and see also Gallagher, 2000), and the growing evidence that the sense of body ownership and the experience of agency over the body rely, at least partially, on distinct mechanisms such as sensory-motor contingencies and cognitive mapping (Arzy and Schacter, 2019; Tsakiris et al., 2007).

### The current work focuses on another conspicuous distinction between processes implicated in the bodily anchoring of consciousness

*error monitoring* (Taylor et al., 2007) on the one hand, and *mirror neuron* activity (Rizzolatti, 2005) on the other. The error monitoring system (EMS) has been hypothesized to continuously track the outcomes of one’s goal-directed actions and detect deviations from the expected performance outcomes. This process, in which the anterior cingulate cortex has been hypothesized to play a key role, is assumed to facilitate adaptive allocation of attentional resources, real-time behavioral adjustments, and learning. The notion that the continuous neural processing of error and prediction errors has a fundamental role in perception and cognition has had a significant impact on theoretical accounts of the mechanisms underlying the illusion of ownership over artificial objects and virtual bodies. Apps and Tsakiris, for example, argued that a sense of ownership over artificial body parts appears as a parsimonious way to reduce conflicts between rich multi-sensory data arriving simultaneously from the biological and the alternative bodies (Apps and Tsakiris, 2014). Ownership illusion is therefore a preferable computational solution, which minimizes the amount of “free-energy” (or surprise) in sensory systems (see Maister et al., 2015).

The mirror neuron system (MNS), on the other hand, is defined by the property that it discharges during both the execution and the observation of specific actions (Keysers and Gazzola, 2010). While the MNS has been extensively studied in the context of social cognition as a mechanism that facilitates embodied communication with *others*, its (right-lateralized) frontoparietal foci have also been implicated in the multimodal experience of the *self*. It has been specifically suggested that the mapping of body-related *perceptual* information onto one’s *motor* system is pivotal for the embodied perception of both others and self (Uddin et al., 2007).

The notion that the EMS and MNS are involved in self-related processing is supported by several neuroimaging studies (for review, see Knyazev, 2013). Of special interest in the context of our work are electroencephalographic (EEG) indices that have been linked with both the theoretical concepts of EMS and MNS on the one hand, and body ownership on the other. First, self-referential processing has been associated with P300 (or P3), which is an extensively studied event-related potential (ERP) component of positive deflection. This ERP emerges between 250-500 ms after unexpected, improbable, and salient stimuli (Polich and Kok, 1995) and is highest over parietal areas. Increased P300 amplitude was observed when participants perceived their own name (Berlad and Pratt, 1995; Fischer et al., 2008; Zhao et al., 2011), face (Ninomiya et al., 1998), and voice (Conde et al., 2015) and in response to self-relevant possessive pronouns (e.g., “my”; Shi et al., 2011; Zhou et al., 2010).

Evidence regarding a specific link between aspects of body ownership and three additional ERP measures has recently accumulated. These measures are positivity on error (Pe), error-related negativity (ERN), and N400. Like P300, Pe is an ERP component that is maximized at parietal recording sites, peaking about 200-500 ms after an erroneous response (Falkenstein et al., 1991; Riesel et al., 2013) across different tasks (Riesel et al., 2013). In fact, Pe is hypothesized to arise from the same source as P300 (Falkenstein et al., 1991; Hester et al., 2005; Nieuwenhuis et al., 2001; Overbeek et al., 2005), and it is suggested to be a special case of P300 that appears in response to infrequent and motivationally-significant errors (Arbel and Donchin, 2011; Overbeek et al., 2005). Pe has been specifically associated with error awareness, unlike the earlier error-related negativity (ERN), which appears also in cases of unaware errors (Di Gregorio et al., 2018; Steinhauser and Yeung, 2010). ERN, which peaks roughly 80 ms after error commission, is largest at frontal and central electrodes (Holroyd and Coles, 2002). It is hypothesized to reflect a key unconscious error-monitoring process (Dehaene, 2018). A third relevant component is N400: a negative potential typically peaking around 400 ms over centro-parietal electrodes after exposure to unexpected semantic events (Kuperberg, 2007). N400 was recently also found to be evoked by perceived unexpected and erroneous actions as well (Balconi and Vitaloni, 2014; Maffongelli et al., 2015).

Two recent studies investigated the relevance of these ERPs to self-perception by introducing involuntary erroneous virtual avatar’s motions. Pavone and colleagues (Pavone et al., 2016) presented faulty virtual object grasping actions from first- and third-person perspectives. They found that perspective and action accuracy interacted so that ERN differences between erroneous and accurate grasping was larger in first-person perspective. Both grasping accuracy and perspective had main effects on parietal Pe, but as the interaction between these factors was insignificant the results indicated that self-perception specifically involves ERN but not Pe response.

In another work by Padrao et al. (Padrao et al., 2016), the sense of agency was temporarily violated as the virtual avatar occasionally performed a movement in an opposite direction to the participant’s motion. Relative to errors made by the participant, forced avatar’s errors elicited a negative parietal N400. This effect correlated with the participants’ self-reported sense of ownership over the virtual body (stronger N400 for larger sense of ownership). Pe was observed after errors, and seemed to be larger following true relative to forced errors, but no statistical test of this difference was reported.

The fourth theoretically-relevant EEG index is Mu suppression. The Mu rhythm (8-13 Hz) is measured at central electrodes, and its suppression assumingly reflects the activation of sensorimotor areas (Fox et al., 2016; Hobson and Bishop, 2017). Mu suppression occurs when an individual either performs or observes motor actions, and is considered as a valid measure of MNS activity (Fox et al., 2016). Whereas most evidence on the involvement of putative mirror-neuron areas in self-recognition comes from functional magnetic resonance imaging studies (see Uddin et al., 2007) one study reported on reduced Mu power during self-related relative to friend-related trait judgments (Mu and Han, 2013).

In addition, three studies demonstrated that Mu is modulated by embodiment manipulations. In two of these studies, body ownership illusion was induced via a synchronous visuotactile stimulation. Evans and Blanke (Evans and Blanke, 2013) reported that the induction of illusion of ownership over a virtual hand evoked reduced suppression relative to a synchronous stimulation of a non-body object or an asynchronous stimulation of both targets. Lenggenhager and colleagues (Lenggenhager et al., 2011) similarly reported on reduced Mu suppression effect when comparing synchronous and asynchronous stroking on the back of the participant’s and an avatar’s back. In a recent work (Shibuya et al., 2018) the authors employed visuotactile stimulation to induce ownership over a real-time video recorded model hand. The synchronous and asynchronous blocks were intersected with single fingers abduction and adduction sequences performed by the model. In this study, Mu suppression during these sequences was found to be larger if they were in the context of synchronous (relative to asynchronous) induction. Mu power also negatively correlated with the self-reported ownership over the virtual hand.

These works powerfully addressed body ownership illusion by inducing *de novo* embodiment vis-à-vis a virtual body rather than eliciting self-related associations that might be confounded by parameters such as familiarity, and affective value. They produce a liminal experience in which an observed avatar’s motion can be perceived either as one’s own or another’s action. Intriguingly, EEG markers of both the EMS and MNS were found to covary with the psychological distance from an agent whose actions were observed (Carp et al., 2009; Kang et al., 2010 for Pe and observational feedback-related negativity respectively, and Aragón et al., 2014 for Mu suppression). Together, the evidence suggests that self-body consciousness and at least some aspects of empathy reside on the same continuum, which implicates both systems.

Thus, the question arises as to the relations between the EMS and MNS in body consciousness. Are these systems similarly affected by the same aspects of embodiment? Are they correlated in their reaction to the induction of body ownership illusion? The small body of research mentioned above does not allow for a straightforward examination of these questions. First, none of these studies tested and compared neurophysiological markers of both systems. Second, a comparison between the studies is difficult since they differ in terms of their design, as the ownership induction phase was not always separated from the probing task.

The present work employs an experimental manipulation, which was designed to activate both the EMS and the MNS. The manipulation included forced spontaneous and instantaneous bounces of a virtual hand following alternative embodiment induction manipulations that were temporally separated from these transient agency violations.

### Our work had two main objectives

(i) Testing the differential impact of parameters that affect body ownership over the EEG markers reviewed above and the correlation between error-monitoring and MNS markers in this context. (ii) Offering EEG markers of body ownership illusion that can be readily integrated into future designs of virtual reality (VR) - a technology that facilitates flexible and controlled body ownership manipulations unachievable by traditional techniques (Blanke et al., 2015).

We specifically examined whether each of the aforementioned EEG markers responds to the following three parameters that were systematically investigated and shown to affect body ownership indices (Ma and Hommel, 2015): (i) The degree to which the participant has agency over the virtual effector (hand); i.e., whether the body ownership illusion was induced by the active participant’s actions or by passive perception of the multisensory stimulation. (ii) The synchronicity of the visuotactile stimulation. (iii) The semantic congruence between the participant’s real hand and the effector; i.e., whether the participant’s hand is virtually represented as a realistic hand-like object.

A complete within-subject factorial design for these parameters would require eight sessions. However, since dozens of stimulus repetitions are often necessary in ERP studies and due to the limited tolerance of participants for prolonged VR use, we adopted a reduced experimental design that tests only main effects of these parameters.

We hypothesized that agency, synchronicity, and semantic congruence will affect each of the four EEG measures following the virtual hand bounces. In each of the tested conditions we further compared the relevant EEG markers with two commonly used measures of body ownership: self-reported ownership over the virtual body (Kalckert and Ehrsson, 2012), and proprioceptive drift towards the location of the virtual body (Botvinick and Cohen, 1998). Finally, we examined whether the EEG effects that are associated with one’s projection onto the virtual body covary with individual tendencies to cognitively and emotionally empathize with others.

## Methods

### Materials

The participants were outfitted with an HTC VIVE headset and four trackers (HTC, New Taipei City, Taiwan), which were placed on their forearms and wrists (Figure 1). The average positional and angular accuracies of the trackers were previously reported to be 1.465 mm and 0.32°, respectively (Yang et al., 2017). The tracking enabled the representation of hand movement relative to the forearm.

**Figure 1:**
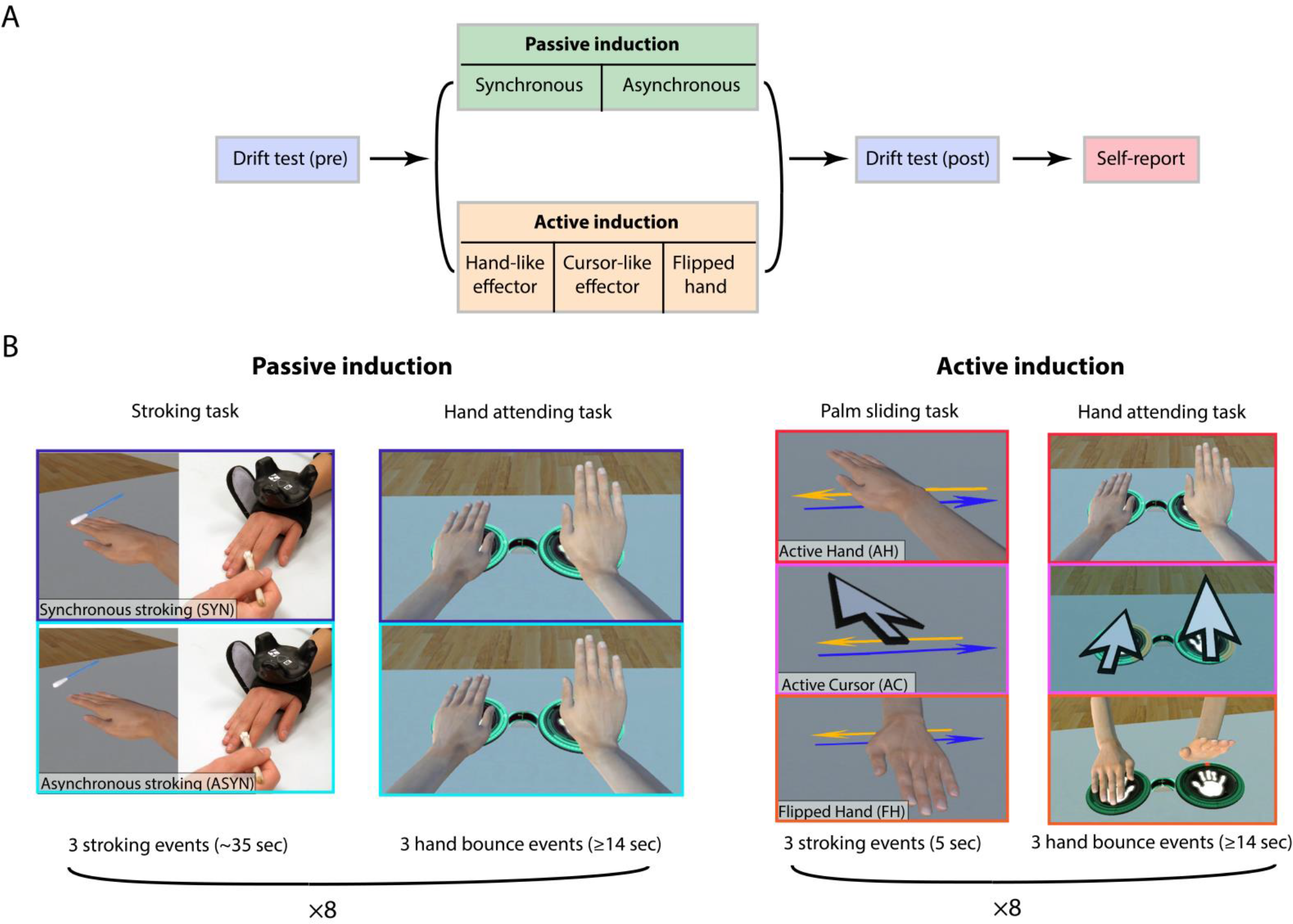
The experimental design. (A) Experiment layout: The participants underwent five sessions, each included different experimental condition (i.e. mode of induction). Before and after the induction of the illusion, proprioceptive drift test was performed. At the end of each session, self-report evaluation was conducted. (B) Modes of induction and *hand attending task* in the different experimental conditions. Left panel: the difference in the stroking task between the passive induction conditions (synchronous and asynchronous) was based on the congruency between the tactile and visual stimulus. Left middle panel - illustration of a hand bounce in the *hand attending task* as appeared in the passive induction conditions. Middle right panel: the difference in the *palm sliding task* between the active induction conditions was based either on hand appearance (*Active Cursor* condition) or orientation (*Flipped Hand* condition). Right panel - *attending task* as appeared in the different active induction conditions.

EEG was obtained using a 16-channel V-Amp amplifier (Brain Products GmbH, Munich, Germany). Passive Ag/AgCl electrodes were positioned on a cap according to the international 10-20 system at 12 recording sites (Fp1, Fp2, Fz, Cz, C3, C4, Pz, P7, P8, Oz, O1 and O2) and referenced online to site FCz. Raw data were recorded using the BrainVision Recorder software at a sampling rate of 250 Hz.

### Participants

Twenty participants were recruited via an advertisement on Facebook. Exclusion criteria were epilepsy, diagnosed attention deficit, and any form of prior head injury or brain surgery. All of the participants had at least 12 years of education and corrected-to-normal vision acuity. All participants signed a consent form approved by the ethical committee of the Tel-Aviv Sourasky Medical Center. One participant was excluded due to poor compliance with experimental procedure. Due to a technical failure the data of another participant was not recorded except for the questionnaires. Thus, valid EEG data were collected from 18 participants (7 females, age: 27.05±2.88 years). Since one of these participants performed the hands’ position estimation incorrectly (centralizing his gaze, but not the headset on the estimated location), he was excluded from the analysis of proprioceptive drift. Interpersonal Reactivity Index data were missing for one participant.

### Procedure

The study outline is presented in Figure 1 and an illustration of the experimental conditions is provided in supplementary video 1. It included four tasks, which were integrated in five experimental sessions. The sessions varied in the method employed to induce or interfere with body-ownership illusion. The participants sat on a chair in front of a real table. During the session, they wore the VR headset, which displayed spatially-aligned virtual table and chair. Their arms and hands were continuously represented in the virtual world. Between sessions, the participants removed the headset and filled out a questionnaire about their sense of body ownership. The tasks were as follows:

1. *Estimation of the real hands’ posit*ion (~6 seconds): The participant was asked to place her hands on two virtual crosses located on the table. Next, the image faded to black and the virtual table reappeared with no representation of the participant’s hands. The participants were asked to point to the estimated location of their *real* left and right hands by placing a virtual circle that was displayed at the center of their field of view. A circular progress bar was activated when the participant had fixated. The estimation was registered after 2 seconds of fixation.
2. *Palm sliding task* (5 seconds): The participant was instructed to track a virtual horizontal arrow that appeared (randomly) either to her left or to her right. She was asked to slide the palm of the hand that was closest to the arrow along this shape for six seconds.
3. *Stroking task* (~35 seconds): The participant was instructed to keep her hands still on the virtual crosses and to keep looking at them. An experimenter stroke her real fingers using a cotton swab either synchronously or asynchronously with an animation of the swab touching the participant’s virtual fingers. The experimenter slid the swab over the participant’s finger. This act was performed either once or three or five times per touching epoch. The number of repetitions was randomized. Three touching trials were included in each block. The target fingers were randomly selected from either the left or the right hand (the side was kept constant during each block).
4. *Hand attending task* (≥14 seconds): The participants were instructed to place their hands on virtual hand symbols located on the table and fixate on a cross located between these symbols. The symbols were encircled with a red progress bar. While the participant watched her virtual hands, brief animations of spontaneous hand bounces were displayed. The animation, which lasted 250 ms, presented the virtual palm bending up to 45° relative to the table and then returning to its initial position. The hand attending block started with 2 seconds of passive viewing and lasted at least 14 seconds depending on the participant’s focus on the fixation cross. Randomly jittered inter-stimulus intervals of 3.55-3.95 seconds separated between the animations. To warrant that the participant’s looks at the hands, the task automatically restarted if the view had not centered on the cross or if the hands were removed from the symbols. Each session included eight blocks per session with at least three hand bounces (and more if the task restarted). To enhance the participant’s attention, a simple detection test was integrated into the task. Once in each block, an object appeared for 200 ms above one of the virtual hands. The shape (sphere, cylinder cube), color (red, green, blue), and side (left, right) of the object were randomized. It was displayed in a random timing starting from 6.5 seconds after the block onset, and at least 1 second after the termination of a spontaneous hand motion. After the progress bar had completed, the participants were asked to briefly answer a recorded question about the shape, color, or side of the object. The average rate of errors was 1.25% per participant (range 0-7.5%).

Prior to the experiment, the participants had a brief initial calibration procedure (aligning the borders and height of the virtual and the actual tables) and had a short (~ 5 min) training session, which introduced each of the tasks. The participants went through five randomly ordered sessions. Each session started and ended with a hands’ position estimation task and included eight hand attending blocks. The sessions differed by the condition of body ownership illusion as follows: (1) *passive synchronous visuo-tactile stimulation* (SYN) included eight blocks of a synchronized stroking task; (2) *passive asynchronous visuo-tactile stimulation* (ASYN) included eight stroking task blocks in which the actual touching either preceded or lagged behind the animated touching (as determined randomly) in two seconds; (3) *Active hand* (AH) included eight palm sliding blocks in which the participant’s hands were represented as virtual hands; (4) *Active cursor* (AC) included eight palm sliding blocks in which the participant’s hands were represented as cursors whose tips were aligned with the tips of the indexing fingers in conditions (1)-(3); (5) Flipped hand (FH) included eight palm sliding blocks in which the participant’s hands were represented as mirror images; i.e., rotated in 180° on the diagonal axis.

In conditions (1)-(3) the virtual arm was elongated so that the palm appeared 20 cm diagonally further from that of the real hand. This shift was also applied to the cursor in condition (4). The position of the virtual crosses was fixed across all tasks and conditions. These conditions allowed for the testing of the three aforementioned parameters that affect embodiment: passivity-activity (SYN-AH), synchronicity (SYN-ASYN), and semantic congruence (AH-AC and AH-FH for violations of morphological and anatomical schemes, respectively).

### Preprocessing

EEG analysis was performed using EEGLAB toolbox (v14.1.1,Delorme and Makeig, 2004)) functions and custom MATLAB scripts. EEG continuous data were first high-pass filtered with a 0.1Hz finite impulse response (FIR) filter and then segmented into epochs according to the task analyzed. Hand attending task epochs started 2 seconds before the spontaneous hand bounce animation and lasted 2 seconds following the bounce. An independent component analysis (ICA) was applied to the epoched data and eye-movement/blink related components were selected and removed by visual inspection.

To mitigate the effect of actual movements on the measured Mu power we discarded trials during which the participants performed significant hand motions. We first computed baseline displacements during all 1.5 seconds epochs that preceded the first bounce in each of the sequences (while the participants were instructed to keep their hands still on the virtual crosses). Trials during which the displacement of the participant’s real hand in any of the three axes exceeded 2.5 standard deviations were excluded. In practice, 8.4% of the trials were discarded. The mean thresholds across all conditions and participants were 3.3, 4.7, and 2.3 mm in the diagonal, vertical, and horizontal axes, respectively.

Time-frequency decomposition was performed on epoched data using the Morlet wavelet transform. Log-spaced frequencies between 5 and 40Hz were estimated using Hanning-tapered window with a linearly increasing width (3 cycles at the lowest frequency and up to 12 cycles). Event-related spectral perturbations (ERSPs, Makeig, 1993) were then calculated by averaging spectral estimates from all trials of each condition and applying baseline normalization (subtracting the mean baseline log power spectrum from each spectral estimate). Since both the right and left virtual hand bounced during the task, ERSPs were averaged in each condition and each subject separately for Central (C3, C4) and Occipital (O1, O2) sites.

For the time–domain ERP analyses, hand attending task epochs were further bandpass filtered between 2–15 Hz, using FIR filters with zero phase distortion, half-amplitude attenuation at the stated frequencies, and transition bands narrower than 1 Hz on either side of the half-amplitude frequency. All epochs were truncated after filtering to the interval −500 to +1000 ms with respect to hand bounce, discarding any edge-effects resulting from the filter window. The filtered epochs were averaged for each individual participant and baseline-corrected to the −500 to 0 ms before the bounce.

## Measures & Analysis

### Self-reported body ownership

We used a Hebrew version of a widely used questionnaire about body ownership illusion (Kalckert and Ehrsson, 2012). The term ‘rubber hand’ in the original questionnaire was replaced with the term ‘virtual hand’. The participants rated the extent to which they agreed with the statements using a 7-point Likert scale ranging from “−3” (strongly disagree) to “+3” (strongly agree). Body ownership scores were computed using the first four items of the questionnaire (e.g., “I felt as if I was looking at my own hand”). Based on previous findings (Ma and Hommel, 2015) we hypothesized that the ownership rating will be larger in the synchronous relative to the asynchronous, the active relative to the passive, and the semantically congruent relative to the semantically incongruent conditions. Therefore, these hypotheses were tested using one-tailed Wilcoxon signed rank test that compared the rating medians. False discovery rate (FDR) correction for dependent tests (Benjamini and Yekutieli, 2001) was applied to control for multiple comparisons.

#### Interpersonal Reactivity Index (IRI)

to assess the participants’ empathic tendencies we used the IRI, which is a standard 28-item self-report measure of trait-empathy (Davis, 1983). This index probes four aspects of empathy: the tendency to transpose oneself into the state of fictitious characters (FS), personal tendencies to adopt another’s point of view (PT), tendency to experience “other-oriented” feelings (EC), and tendency to feel distressed when faced with another’s distress (PD). In line with a common practice in the field of empathy research (e.g., Atoui et al., 2018; Bock and Hosser, 2014; Harari et al., 2010; Hooker et al., 2010; Kerr-Gaffney et al., 2019) we combined FS and PT into the “cognitive empathy” factor and EC and PD into the “affective empathy” factor.

#### Proprioceptive drift

The drift was calculated as follows:

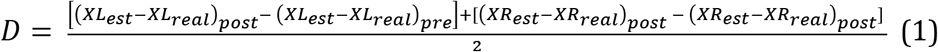

where XL_est_ and XR_est_ are the diagonal coordinates of the self-estimated locations of the participants’ left and right indexing finger respectively, and XL_real_ and XR_real_ refer to the real location averaged over the fixation period (two seconds) during which the estimation was performed. Since Kolmogorov–Smirnov test rejected the hypothesis that the drift data were normally distributed, we used FDR-corrected non-parametric two-sided Wilcoxon signed rank tests to examine whether the drift in each of the conditions differed from zero. Since previously (Ma and Hommel, 2015) not all of the three factors (synchronicity, semantic congruence, and agency) had main effect on proprioceptive drift, we conducted two-sided Wilcoxon signed rank tests for hypotheses about differences between the conditions SYN-ASYN, SYN-AH, AH-AC, and AH-FH.

#### Hand motion following virtual hand bounces

To examine whether the induction condition affected the participants’ actual hand movements following the virtual hand bounce, we registered the coordinates of the real hands within an interval of 1500 ms after the bounce onset. The range of the movement within each of the axes in this interval served as a dependent measure in this analysis. Since a Kolmogorov–Smirnov test indicated that the data is not distributed normally, we used statistical tests that do not assume normality. First, we examined whether the axis has a main effect on the movement using two-way ANOVA with the factors AXIS (x,y,z) and CONDITION (synchronous, asynchronous, active hand, active cursor, and flipped hand) and permutation tests to estimate the significance of the results. We permuted the AXIS 10,000 times and defined the p value as the number of permuted cases in which a larger main effect was obtained divided by the total iteration number. Pairwise differences between the conditions were tested using Wilcoxon signed rank tests.

### Event-related spectral perturbation (ERSP) analysis

To identify a specific time-frequency window in which the conditions differed the most we used a non-parametric cluster based permutation test (Maris and Oostenveld, 2007), which essentially controls the false alarm rate when running multiple comparisons on these kind of data. Each time-frequency bin was subjected to a one-way repeated measures ANOVA with 5 conditions, from which significant bins (p<0.05) were further clustered and tested using a permutation test. For this analysis, we averaged the power of each frequency band at each time window in the cluster that showed a CONDITION main effect. We first tested whether the power in the observed cluster was below baseline using FDR-corrected two-sided signed rank test for each of the conditions. For post-hoc analyses of the main effects of agency, synchronicity, and semantic congruence we conducted paired Student’s t-tests comparing the mean power in the cluster for the following contrasts: SYN-ASYN, SYN-AH, AH-AC, AH-FH. Permutation tests were performed to estimate the significance of the results. In these tests, the t statistic of the original comparison was compared with the t distribution generated by 10,000 identical tests that compared couples of data series in which the two relevant condition labels were randomly shuffled. The two-tailed p value was defined as the proportion of t values in the background distribution that were equal or greater than the original. FDR correction was applied to control for the four comparisons. The power at the observed cluster was also compared to zero for each of the conditions using a two-tailed signed rank test.

### Error-related ERPs

Similarly to the ERSP analysis, separate one way ANOVAs were conducted on the event-related average time-courses of midline electrodes (Fz, Cz, Pz and Oz) in order to identify time windows of significant difference between the conditions. Cluster-based permutations were used to correct for multiple comparisons in the time domain and post-hoc analyses were performed to examine the main effects of the independent variables.

### Relationships between the behavioral and EEG measures

To examine the relationship between the measures, we conducted pairwise comparisons between the self-reported measures, proprioceptive drift, and the EEG indices that showed significant effects in previous tests. We compared the variables using Pearson correlation, but since a Kolmogorov–Smirnov test indicated that they do not distribute normally, we estimated the significance of the results using a permutation test as recommended in such cases (e.g., Bishara and Hittner, 2012). We randomly redefined pairs of EEG and drift variables, repeating this procedure 10,000 times. The two-tailed probability value was defined as the proportion of the coefficients obtained in the permutation tests that were equal or greater than the original coefficient in absolute value. Spearman’s tests were used to compare the EEG indices with ordinal ownership rating, and with the cognitive and affective empathy scores using the same permutation method. The correlation between the proprioceptive drift and the ownership rating was examined using the same tests. FDR correction was applied within each set of five pairwise comparisons between the measures.

#### Data and code availability

Data and code are available upon request from the authors under the institutional confidentiality regulations.

## Results

### *Behavioral measures* (Figure 2)

As expected, body ownership scores (Figure 2a) were significantly higher in the synchronous relative to the asynchronous conditions (Z=1.94, one-tailed p=0.03, Q_FDR_=0.05), and in the active relative to the flipped hand condition (Z=2.28, p=0.02; Q_FDR_=0.05). No significant difference was found between the active and the passive conditions (Z=−1.58, p=0.94) and the active hand and cursor (Z=1.42, p=0.08).

**Figure 2:**
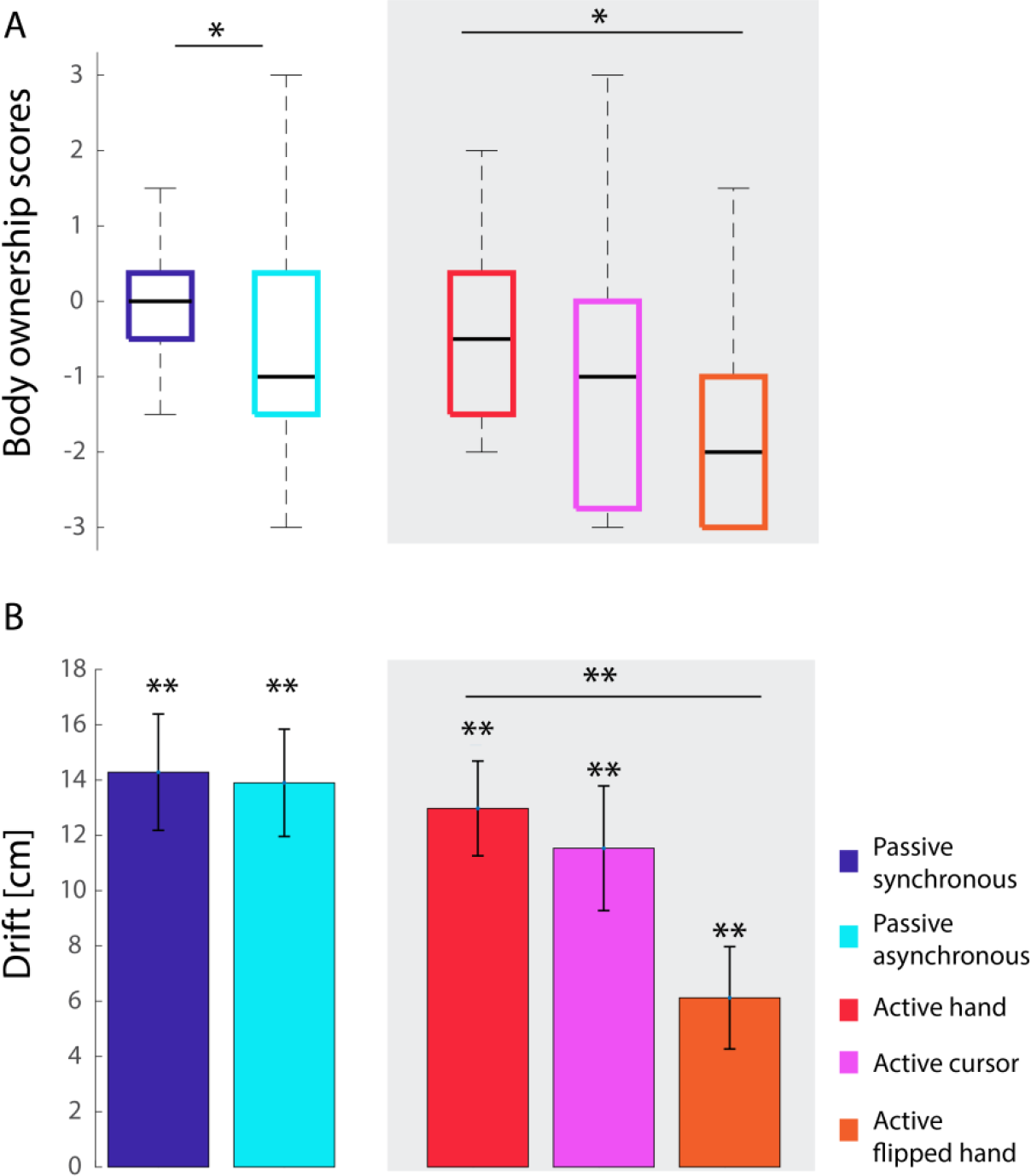
The impact of embodiment manipulations on behavioral measures. A. Box plots of self-reported body ownership scores. The central marks represent the medians and the edges of the box indicate the 25th and 75th percentiles. B. Proprioceptive drift. * QFDR<=0.05, ** QFDR<=0.001.

### A significant proprioceptive drift (Figure 2b) was evident in all conditions

SYN (Z=3.57, p=0.0003, Q_FDR_=0.0004), ASYN (Z=3.62, p=0.0003, Q_FDR_=0.0004), AH (Z=3.62, p=0.0003, Q_FDR_=0.0004), AC (Z=3.62, p=0.0003, Q_FDR_=0.0004), and FH (Z=2.67, p=0.008, Q_FDR_=0.008). The drift was significantly higher for AH relative to FH (Z=3.53, p=0.0004, Q_FDR_=0.002). Conversely, no significant differences were found between SYN and ASYN (Z=0.21, p=0.83), SYN and AH (Z=0.4, p=0.68), and AH and AC (Z=0.97, p=0.33).

Significant correlation between the ownership scores and proprioceptive drift was found in the case of SYN (R_Spearman_(15)=0.59, p=0.01; Q_FDR_=0.05). Marginally significant correlation was found in AC (R(15)=0.49, p=0.02; Q_FDR_=0.06). In the three other conditions, the correlation was not significant (R(15)=0.17, p=0.24; R(15)=0.25, p=0.16; R(15)=−0.06, p=0.41 in ASYN, AH, and FH, respectively). The proprioceptive drift was marginally correlated with the affective empathy scores in SYN (R_Spearman_(15)=0.52, p=0.02; Q_FDR_=0.06), ASYN (R(15)=0.5, p=0.03; Q_FDR_=0.06), and AH (R(15)=0.41, p=0.05; Q_FDR_=0.09), but not in AC and FH (R(15)=0.25, p=0.17; R(15)=0.29, p=0.14, respectively). Affective empathy was not correlated with the ownership scores (R_Spearman_(16)=0.1, p=0.7; R(16)=0.27, p=0.65; R(16)=0.02, p=0.95, R(16)=−0.12, p=0.3; R(16)=−0.01, p=0.96 for SYN, ASYN, AH, AC, and FH, respectively). Cognitive empathy correlated with neither the drift (R_Spearman_(15)=−0.15, p=0.59; R(15)=−0.37, p=0.15; R(15)=0.22, p=0.4, R(15)=0.02, p=0.94; R(15)=−0.18, p=0.55) nor the ownership scores (R_Spearman_(16)=−0.02, p=0.94; R(16)=0.28, p=0.29; R(16)=0.35, p=0.16, R(16)=0.05, p=0.85; R(16)=0.29, p=0.26).

### Hand motion following virtual hand bounces

When testing the actual movement of the participants’ hands following the virtual hand bounce, we found a main effect of AXIS (F(2,255)=17.64, p<0.0001). Movement on the vertical axis (corresponding with the direction of the virtual hand’s movement) was significantly larger than on the horizontal (t(17)=2.86, p=0.0009) and the diagonal (t(17)=3.23, p=0.0002) axes. The average range of movement (across conditions) was 0.71±0.17, 1.08±0.9, and 0.66±0.15 mm in the horizontal, vertical, and diagonal axes, respectively. However, when comparing pairs of conditions for physical hand movement on the vertical axis, no significant difference was found (Z=0.02, p=0.98; Z=0.89, p=0.89; Z=0.94, p=0.34; Z=−1.6, p=0.1 for SYN-ASYN, SYN-AH, AH-AC, and AH-FH, respectively).

### Mu suppression

A main effect of CONDITION (p(corrected)=0.035) was observed in a cluster, which largely falls within the Mu band between 95-824 ms after the onset of the bounce (Figure 3B). No similar significant condition effect was found for occipital (min p=0.108) electrodes. Figure 3A presents the effects in each of the conditions. The power at the observed cluster was significantly lower than baseline in the test conditions SYN (Z=−2.5, p=0.01, Q_FDR_=0.03) and AH (Z=−3.2, p=0.001, Q_FDR_=0.006), but not in the control conditions ASYN (Z=0.15, p=0.88), AC (Z=−0.06, p=0.95), and FH (Z=−0.24, p=0.81).

**Figure 3:**
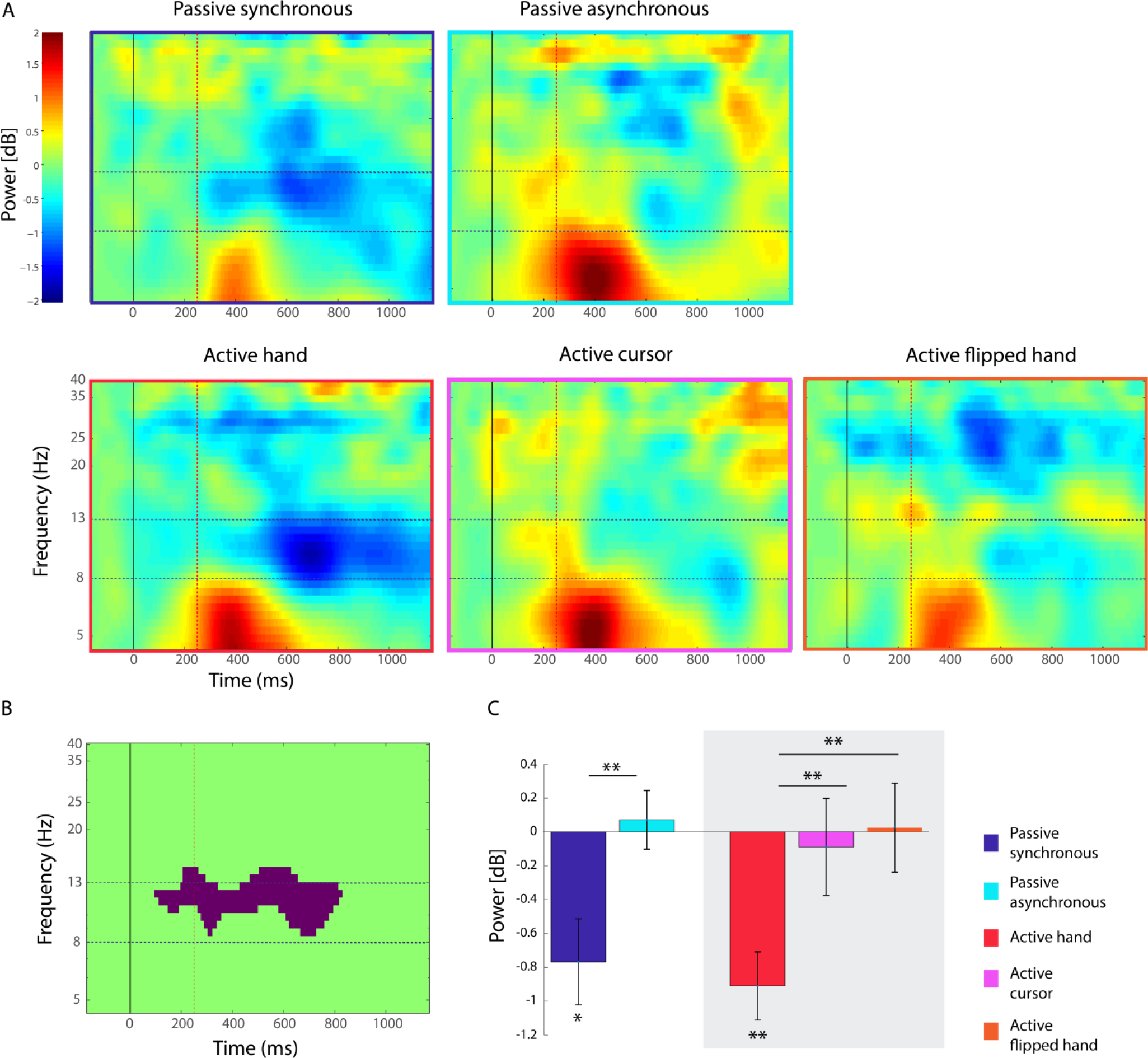
Mu power is modulated by body ownership manipulations. (A) Event-related spectral perturbations in the five conditions. The horizontal dashed lines indicate the boundaries of the Mu band. The vertical black and red lines indicate the onset and end of the virtual hand bounce animation, respectively. (B) The results of one-way ANOVA of the ERSPs over conditions. The purple area represents a cluster that survived correction for multiple comparisons. (C) mean power of the cluster demarcated in (B) for each of the conditions. The vertical lines indicate ±1 standard error. * Q_FDR_<=0.05, ** Q_FDR_<=0.01.

Post-hoc analyses indicated greater power reduction in this Mu-dominated cluster (Figure 3C) in SYN relative to ASYN (t(17)=3.21, p=0.005, Q_FDR_=0.01, Cohen’s d=0.94), AH relative to AC (t(17)=2.77, p=0.01, Q_FDR_=0.02, d=0.8), and AH relative to FH(t(17)=4.79, p=0.0003, Q_FDR_=0.001, d=0.97). No significant difference was found between the two test conditions SYN and AH (t(17)=0.50, p=0.61).

### Error-related ERPs

A main effect of CONDITION was evident only in electrode Pz at a time window 328-380 ms after the bounce onset (p(corrected)=0.03). This positive deflection corresponds to the Pe/P300 wave. In the main effect analyses (Figure 4B), Pe/300 was found to be lower in AH relative to AC (t(17)=2.87, p=0.01, QFDR=0.02, d=0.44) and FH (t(17)=3.04, p=0.01, Q_FDR_=0.02, d=0.41). No significant difference was found between SYN and ASYN (t(17)=0.8, p=0.45) and SYN and AH (t(17)=0.69, p=0.5).

**Figure 4:**
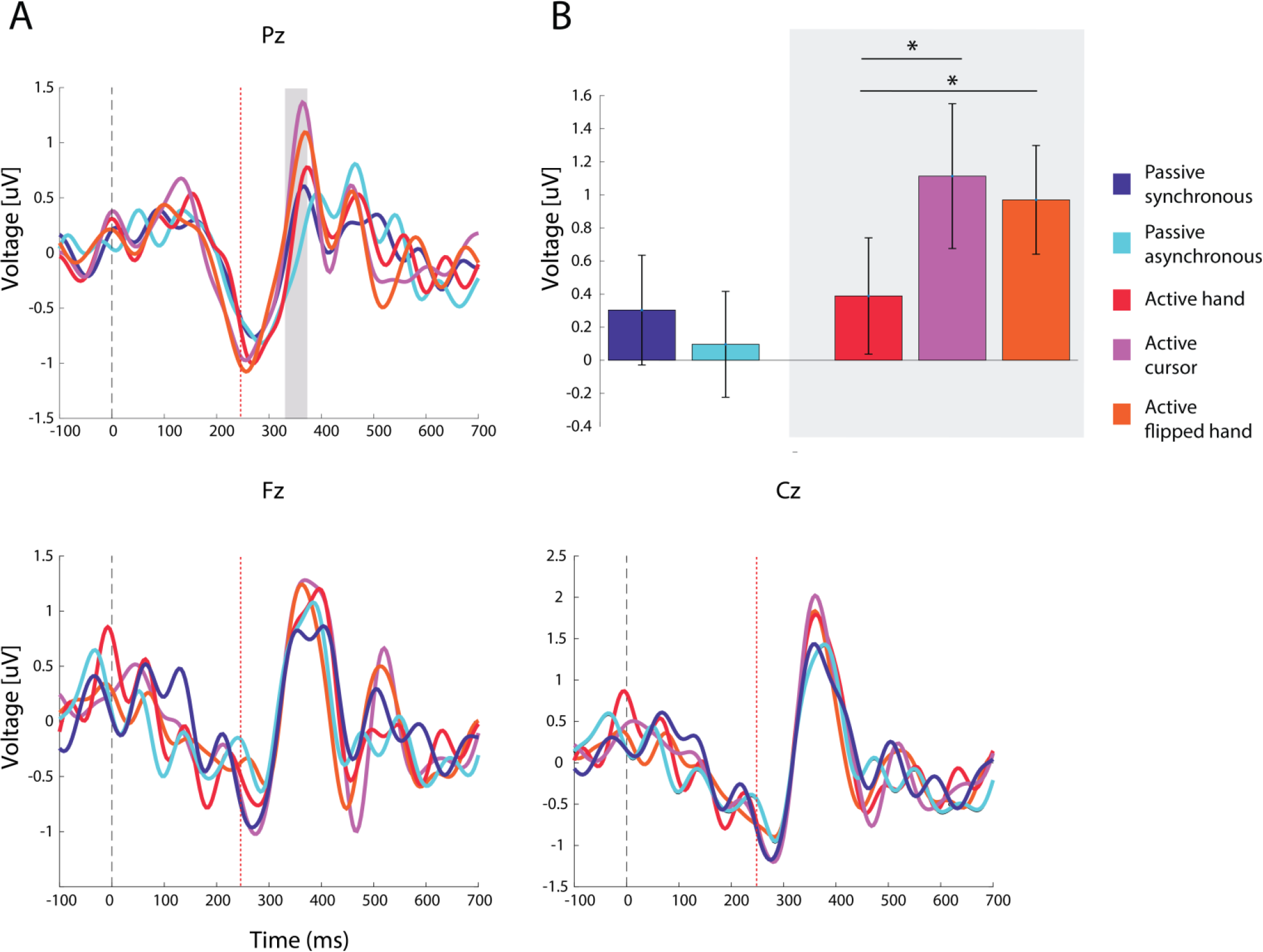
ERPs following virtual hand bounces. (A) response-locked average time-courses at frontal (Fz), central (Cz) and parietal (Pz) electrodes. The vertical black and red lines indicate the onset and end of the virtual hand bounce animation, respectively. The gray surface frames a time window in which one-way ANOVA showed a main effect of CONDITION (corrected p<0.05). (B) Mean amplitudes at the time window in which a significant main effect of CONDITION was observed. The vertical lines indicate ±1 standard error. * Q_FDR_<0.05.

### Relationships between the behavioral and neural measures

The correlations between the neural, behavioral, and self-reported measures are presented in Table 1. Marginally significant negative correlation was found between the Mu power and the ownership scores in the case of ASYN (R_Spearman_(16)=−0.54, p=0.01. Q_FDR_=0.06). The Mu power correlated with neither the questionnaire scores nor the proprioceptive drift.

**Table 1:**
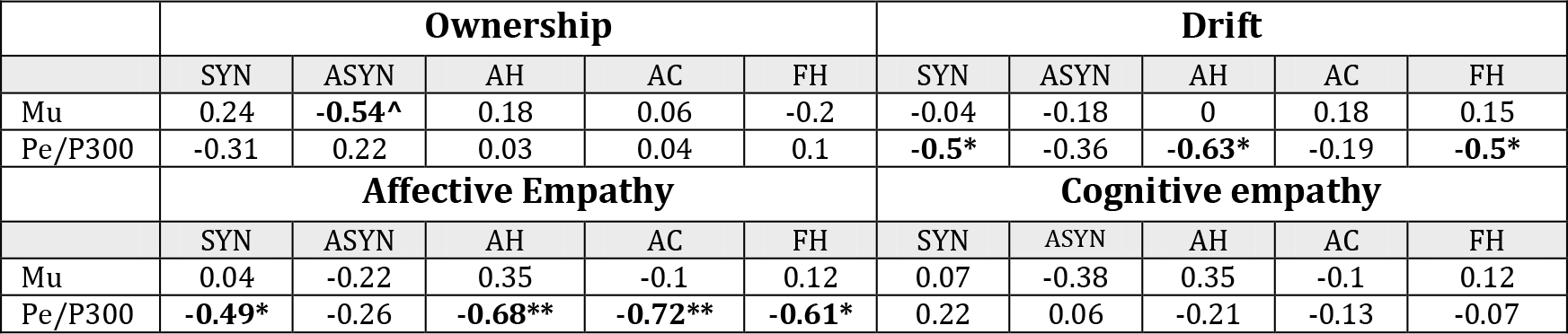
relations between the neural (Pe/P300 amplitude and Mu power), behavioral, and self-reported measures. The values in the table are Spearman coefficients, except in the case of the drift analysis in which Pearson coefficients are presented. ^Q_FDR_=0.06, * Q_FDR_<0.05, ** Q_FDR_<0.01.

The Pe/P300 amplitude was not significantly correlated with body ownership scores. However, the proprioceptive drift and the Pe/P300 magnitude were negatively correlated in the conditions SYN (r(15)=−0.5, p=0.02. Q_FDR_=0.03), AH (r(15)=−0.63, p=0.004, Q_FDR_=0.02), and FH (r(15)=−0.5, p=0.02, Q_FDR_=0.03), but not in ASYN (r(15)=−0.36, p=0.08) and AC (r(15)=−0.19, p=0.23). Furthermore, Pe/P300 amplitude negatively correlated with the affective empathy scores in AH (r(15)=−0.68, p=0.001, Q_FDR_=0.003), SYN (r(15)=−0.49, p=0.02, Q_FDR_=0.03), AC (r(15)=−0.72, p=0.001, Q_FDR_=0.003), and FH (r(15)=−0.61, p=0.006, Q_FDR_=0.01). No significant correlation was found for ASYN (r(15)=−0.26). The cognitive empathy scores were not significantly correlated with Pe/P300 in any of the cases.

Finally, no significant correlation was found between Mu power and Pe/P300 amplitude in any of the conditions (r(16)=−0.27, p=0.28; r(16)=0.09, p=0.74; r(16)=0.06, p=0.81; r(16)=−0.22, p=0.38; r(16)=−0.01, p=0.95 for SYN, ASYN, AH, AC, and FH, respectively).

## Discussion

Employing experimental stimuli that target both the MNS and EMS, our work examined how specific EEG indices are affected by three parameters of body ownership: visuotactile synchronicity, semantic congruence, and agency. The findings indicate that two of the tested EEG indices are indeed sensitive to these parameters: Mu power and Pe/P300. However, in line with the notion that the responses of these EEG indices to the avatar’s spontaneous motions represent distinct aspect of body ownership, they showed different patterns of covariance with the tested variables.

First, Mu power and Pe/P300 were not correlated across participants in any of the conditions. Second, these indices differed in their sensitivity to parameters of body ownership manipulation. In response to the virtual hand bounces, Mu power was modulated by both synchronicity (SYN-ASYN) and semantic congruence (AH-AC, AH-FH), whereas Pe/P300 was modulated only by semantic congruence (AH-AC, AH-FH). Moreover, in both test conditions (SYN, AH), as well as in other control conditions, Mu power and Pe/P300 significantly correlated with the proprioceptive drift and the affective empathy scores, whereas Mu power correlated with neither of them. Finally, body ownership manipulations influenced Pe/P300 and Mu power in opposite directions. While Mu suppression was *enhanced* with synchronous visuotactile simulation and semantic congruence, the Pe/P300 effect was *attenuated* by the latter parameter. Moreover, in the test conditions Pe/P300 amplitude was decreased with higher proprioceptive drift and affective empathy scores. These findings point to an intriguing dissociation between the effects of spontaneous hand bounces on Pe/P300 and Mu suppression in terms of the functions they represent.

### Mu suppression as an indication for enhanced sensorimotor sensitivity

The interpretation of the effects of our experimental manipulations on the Mu power is relatively straightforward. As the average measured vertical displacement of the participant’s real hand was below detection threshold and trials with extreme movements (<2.5 std) were discarded from analysis, it is reasonable to interpret the Mu modulations by SYN and AH as reflecting mainly motor activations by visual input. To restate Shibuya et al.’s interpretation of their similar findings (Shibuya et al., 2018), the enhanced Mu suppression in SYN and AH possibly reflects increased back projection of sensory information about self-motion. As mentioned above, evidence suggests that the MNS encodes not only *motor* information about *own* motions and *perceptual* information about *others’* motions, but also *perceptual* information about *own* motions. The increased Mu suppression in SYN and AH may therefore reflect an enduring enhancement of sensory feedback in the MNS due to the participant’s increased tendency to perceive the virtual hands as her own hands.

Our findings join to recent evidence on the link between synchronicity and *enhanced* Mu suppression (Shibuya et al., 2018). At the same time, they seemingly conflict with previous findings on *reduced* Mu suppression under synchronous visuotactile stimulation (Evans and Blanke, 2013; Lenggenhager et al., 2011). However, a key difference between the studies should be emphasized. The reported reduced Mu suppression was found *during* the synchronous relative to the asynchronous stroking and it may reflect greater multisensory visuo-tactile conflict under this condition (Evans and Blanke, 2013), whereas the increased Mu suppression is Shibuya et al.s’ work was found *after* the synchronous stroking, possibly reflecting induced enhancement of back projection of the MNS to sensory circuits (Shibuya et al., 2018).

These findings can therefore be seen as a replication of Shibuya et al.’s observations about the effects of synchronous visuotactile stimulation over Mu power vis-à-vis spontaneous movements in a fully virtual environment. Our findings further suggest that the Mu power may be induced not only by synchronicity of bottom-up tactile and visual cueing, but also by the congruence of top-down presentations.

In regard to the synchronicity effect, it should be emphasized that since the induction phase (stroking) was temporally separated from the probe phase (bouncing hands), the enhanced suppression is unlikely to reflect an increased attentional focus on the correlated compared to the uncorrelated multisensory stimulation. Such attentional effect, if exists, would probably be limited to a shorter time window (Press et al., 2007), whereas the observed Mu suppression enhancement indicates a persisting effect.

Admittedly, our behavioral results do not shed light on the functional significance of the observed Mu effect, since it was only marginally correlated with ownership scores and only in the ASYN control condition. It seems that the link between the Mu effect and the sense of ownership should we examined using probes of other behavioral aspects of embodiment that are not reflected by the ownership questionnaire and the proprioceptive drift. Specifically, the interpretation outlined above hypothesizes a link between the Mu effect and the aspect of perceptual sensitivity to the virtual body. Future studies may explore this link by adapting sensory gating measures such as prepulse inhibition (Bastin et al., 2017) to virtual setup.

### Pe amplitude reduction and error buffering

While the enhancement of Mu suppression in SYN and AH is in line with expectations about the enhanced involvement of the MNS in these conditions, the reduced Pe/P300 amplitude in AH relative to AC and FH and its negative correlation with the proprioceptive drift and affective empathy scores are less intuitive. What may explain the observations that Pe/P300 amplitude was *reduced* in a condition of increased semantic congruence and that it also tends to decrease more among participants who manifest higher perceptual drift toward the virtual body? Why was this component modulated by semantic congruence but not by synchronicity?

A recent mechanistic account of Pe offers relevant insights in this context. While Pe has been consistently associated with awareness to errors (e.g., Endrass et al., 2007; Hewig et al., 2011; Hughes and Yeung, 2011) and response confidence (Boldt and Yeung, 2015; Hewig et al., 2011), Steinhauser and colleagues (Steinhauser and Yeung, 2012, 2010; Steinhauser et al., 2018) have suggested that Pe reflects not awareness or confidence *per se*, but rather an underlying decision process in which evidence for response error or response conflict is accumulated until it reaches a certain threshold. Using computer simulations and experimental manipulations, these researchers demonstrated an association between the strength of evidence for an error on the one hand and Pe amplitude on the other. Steinhauser and his colleagues employed a visual detection task where the participants were asked not only to perform accurately, but also to provide a post-trial feedback on the accuracy of their performance. In one study (Steinhauser and Yeung, 2010), the researchers used two alternative reward schemes that encouraged more or less frequent error signalling and assumingly induced the assignment of lower and higher weights for evidence that an error has occurred, respectively. In a later study (Steinhauser and Yeung, 2012), the researchers manipulated the speed-accuracy trade-off, assuming that an evidence for an error has higher weights under high speed pressure. In both studies, Pe amplitude after response to the primary task was found to be lower under conditions of weaker evidence for an error.

Importantly, this group has recently extended its notion of Pe as reflecting evidence accumulation beyond perception to the domain of action (Steinhauser et al., 2018). Steinhauser and colleagues demonstrated that unexpected action outcomes in a primary task (puzzle solving) is followed by an increase in Pe amplitude after errors in a subsequent task (color flanker). They conclude that the unexpected action effect in the first task increased the sensitivity to subsequent errors, thus indicating the existence of a link between various error monitoring processes. The authors posit that the common error monitoring system accumulates and integrates evidence of error from cognitive, sensory, and autonomous sources.

The notion that Pe reflects the accumulation of evidence on errors and unexpected action outcomes may explain the direction of the Pe effect observed in our study. The semantic incongruence in the AC and FH conditions functions as a persistent source of top-down conflict, which possibly accumulates with the transient unexpected virtual hand bounces to elicit enhanced Pe effects. This semantic conflict is mitigated in the AH condition, which involves higher similarity between the real participant’s hand and its virtual representation. In keeping with Steinhauser et al.’s account, the decreased Pe amplitude in AH may therefore reflect a reduced load of conflicting evidence on the EMS. Afterall, the tolerance of the brain to violations of cross-modal expectations under a certain threshold of magnitude has been hypothesized to facilitate the perceptual flexibility needed for a body ownership illusion (see Blanke et al., 2015; Maister et al., 2015).

The observation that ASYN did not involve higher Pe effect compared to SYN (the Pe amplitude in both conditions was low, as in AH), indicates that the synchronicity of low-level perceptual processes has little impact on this ERP wave. Alternatively, it is possible that the key difference between the SYN-ASYN contrast on the one hand and AH-AC and AH-FH contrasts on the other was the lasting conflict in the latter case. In these conditions, the semantically incongruent hand representations were active throughout both the priming (palm sliding) and probe (hand attending) tasks, whereas in ASYN the perceptual synchronicity violation was limited to the priming task (stroking). Future research may examine the differential contribution of the type of manipulation and its duration on Mu and Pe/P300 effects.

The impact of bottom-up manipulations on these measures may be gradual. For example, the Pe/P300 effect may decrease with the resemblance between the avatar and the participant (e.g., in terms of skin color). In light of intriguing evidence that body illusions in VR facilitate the adoption of perceptual and conceptual aspects that are related to an avatar and may even reduce implicit racial bias (e.g., Banakou et al., 2013; Hasler et al., 2017; Peck et al., 2013), future work could examine whether the Mu and Pe/P300 effects observed in our study correlate with these behavioral changes. In this context, it is worth highlighting the marginal correlation between the drift and the affective empathy scores that was found in three of the conditions, and the correlation between the affective scores and Pe/P300 amplitude in four of the conditions. These findings call for a further investigation of the relation between these parameters as the tendencies to share the affective perspective of another person and to adopt an illusion of ownership on a virtual body may be linked. It is possible that Pe/P300 reflects tolerance to challenges to these attachment processes as an index of immersion.

### Caveats and limitations

Finally, several issues and caveats of the current study should be noted. First, the two measures of ownership - the questionnaire scores and the proprioceptive drift - were not consistently correlated in our study across conditions. However, the large body of evidence on the relation between these measures is in itself inconclusive: while some studies reported on significant correlations between drift and ownership rating (e.g Botvinick and Cohen, 1998; Kammers et al., 2009; Tsakiris and Haggard, 2005). other found no correlation (e.g. Abdulkarim and Ehrsson, 2016; Holle et al., 2011; Matsumiya, 2019; Riemer et al., 2015) (see Shibuya et al., 2017 for a similar discussion). Our findings point to the possibility that the relations between these measures depends on the induction modes, as they may be most correlated under passive synchronous stroking. Further systematic research is required to establish this link.

Second, the difference in ownership rating between SYN and ASYN in our study was minor, and at odds with our expectations no significant rating difference was observed between AH and AC. Furthermore, while all conditions involved a significant proprioceptive drift, no significant drift differences were evident when comparing SYN and ASYN or AH and AC.

Reports on proprioceptive drift following asynchronous visuotactile stimulation are not common but also not unprecedented (see Rohde et al., 2011; Tsakiris and Haggard, 2005). Some features of our specific design possibly enhanced it. Apparently, the asynchronous condition in our study was not entirely devoid of multisensory synchronicity. During the initial calibration and at the beginning of the block, the participant stretched her hand and touched the table while receiving visual input that was matched with tactile and proprioceptive information. This minor synchronous stimulation might have compromised the difference between SYN and ASYN. Thus, as a methodological improvement in future studies, the display of the hand representations could be limited to the primary and probe tasks.

In regard to the insignificant differences in the behavioral measures between AH and AC, it is worth mentioning a work by Ma and Hommel’s (Ma and Hommel, 2015) that similarly found no proprioceptive drift difference between ownership induction conditions in which semantically congruent and incongruent objects were used. Importantly, the authors reported that the drift toward an embodied virtual rectangle was evident under a condition in which the participant actively manipulated this object that represented her hand. They make the claim that the active induction may impose a bottom-up agency that overrides the top-down semantic incongruence and induces a sense of ownership despite semantic incongruence. Thus, it is possible that the active agency in the AC condition in our study “masked” the behavioral effect (but interestingly, not the Mu and Pe/P300 effects).

An additional factor that might have compromised this effect is the selection of the cursor shape as a non-body object. A previous work that similarly induced a sense of ownership over a virtual cursor reported that this condition was not consistently scored lower than a virtual hand condition in questions about immersion and ownership (Yuan and Steed, 2010). It is possible that since cursors are commonly used as indicators of the user’s agency on computer screens, this specific shape is not the optimal exemplar for semantically incongruent representation of the participant’s hand. Future studies may therefore adopt alternative semantically incongruent hand representations.

### Methodological and scientific significance

Our observation that the Pe/P300 and Mu effects are sensitive to body ownership manipulations suggests that they can be used to probe this process especially under passive synchronous visuotactile stimulation and active movement of realistic virtual body representations. Unlike measures such as proprioceptive drift, self-reported ownership, and galvanic skin conductance response to threat on the virtual body, which are usually applied once per experimental block or as pre-post difference measures, the virtual hand bounces can be spread throughout the experimental design as transient cues. Thus, the continuous measurement of these neural markers potentially enriches the arsenal of probes employed in the emerging field of virtual body ownership illusions.

Beyond basic science, the validation of Pe/P300 and Mu suppression as body ownership markers may be instrumental in diagnostic and rehabilitation contexts. Rehabilitation protocols based on VR have been recently developed for various motor dysfunctions, including post-stroke deficits (Faria et al., 2018; Laver et al., 2017). VR is an attractive rehabilitation medium since it provides a fully controlled environment, which allows for an iterative treatment and an immediate feedback and reinforcement on the patient’s performance. The hand bouncing manipulation and the EEG markers discussed above can be readily integrated into such rehabilitation protocols, allowing continuous monitoring, feedback, and reinforcement. This may be particularly valuable in VR rehabilitation of patients with impairment in basic movement capabilities. A recent work (Ossmy and Mukamel, 2016) pointed to the potential of VR in such cases by showing that motor training of the right hand, which was displayed in real-time as movements of the immobile left hand, can enhance the performance of the latter.

## Conclusions

To sum up, our findings demonstrate that two of the tested EEG components show distinct patterns of reaction to spontaneous involuntary movements of embodied objects. Mu suppression, which is linked to the MNS, increases with both semantic congruence and synchronicity of the induction. Pe/P300 amplitude, on the other hand, decreases with semantic congruence, all the more so in participants who showed higher proprioceptive drift in AH and manifest higher affective empathy tendencies. These findings suggest that while synchronicity and semantic congruence induce a lasting sensorimotor sensitivity to the alternative body, the latter parameter also reduces the sensitivity of the EMS to minor erroneous motions of that body. The observed effects also have methodological and scientific potential as MNS and EMS markers, respectively, in body ownership illusion in VR.

## Acknowledgments

We would like to thank Gil Asher, Sraya Harif, Shani Libi, and Matanel Libi from Actiview Ltd for producing the VR content. We also thank Yoav Zamir for critical technical assistance, and Talma Hendler for a valuable support. NG was supported by the NARSAD Young Investigator Award provided by the Brain & Behavior Research Foundation and the Oxley Foundation, and GG was supported by the Sagol family fund.

